# An analysis of natural variation in *Pinus pinaster* through the lens of systems biology

**DOI:** 10.1101/2024.01.29.577781

**Authors:** Jesús Pascual, Cristina López-Hidalgo, Isabel Feito, Juan Majada, Mónica Meijón

## Abstract

*Pinus pinaster* is a main species in Mediterranean forests, being naturally distributed through the Mediterranean basin, covering regions with a variety of geoclimatic conditions. This distribution in different environments leads, by natural selection, to a natural variation within the species that manifests at phenotypic level in populations with different growth features and overall tree architectures. Studying *P. pinaster* natural variation is necessary to understand the genetic heritage of the species and can provide valuable information for information-based decision-making regarding forest management and breeding programmes. In this paper, we analyzed the natural variation in needles and buds from three provenances from contrasting geoclimatic locations using a common garden approach and proteomics. The integration of the proteomics data with tree growth-related parameters, geoclimatic features at provenances original locations, and sample-matched metabolomics data previously generated provided novel knowledge on metabolism rearrangements related to secondary metabolism and associated to growth features and the adaptation to light and UV-B radiation intensities.

## Introduction

Maritime pine (*Pinus pinaster* Aiton) is a widespread medium-size tree native to the western Mediterranean basin. In fact, it is one of the most common tree species in the Mediterranean area (Viñas et al., 2016). It stands out for its rapid growth and undemanding behavior, and it is a central species in natural forests and forestry for its economical added value. Due to these characteristics, it is widely used in soil protection, reforestation and intensive plantations aimed at wood production. Its wood is appreciated for producing construction wood and furniture and it represents an important part of the existing timber stock (Carle, 2014). Thus, most *P. pinaster* distribution corresponds to plantations, being considered as a highly invasive species in the southern hemisphere, where it has been introduced for environmental and economical purposes. Natural populations are distributed throughout the Mediterranean basin. These populations are classified in six major groups according to contrasting geoclimatic conditions across the natural geographic distribution of the species: Continental France, Corsica, Northwestern Spain, Southeastern Spain, Morocco and Tunisia (Bucci et al., 2007). Such a distribution in locations with different environmental conditions in terms of temperature, rainfall, irradiance or soil composition leads to different natural selection pressures that derive in the selection of different genetic variants in each population (Cañas et al., 2015). This genetic variation, associated with a natural and artificial distribution in a rather wide range of geographical locations, results in populations with a phenotypic variation and differential traits regarding, for example, stress susceptibility (L Corcuera et al., 2012; Gaspar et al., 2013; Öquist and Huner, 2003; Yakovlev et al., 2012). Altogether, these populations represent the natural variation of the species. Studying natural variation can help to decipher adaptation mechanisms and to know and understand the genetic heritage represented by a species, what can be useful for knowledge-based decision-making and natural resources management. It can also help to understand the processes governing tree development and determining tree architecture, relevant from an industrial point of view.

In *P. pinaster*, natural variation is reflected in molecular biomarkers and quantitative traits (L Corcuera et al., 2012; González-Martínez et al., 2002). Maritime pine natural variation has been also studied at metabolome level using a common garden approach that allowed to uncover differences at secondary metabolism, especially regarding flavonoids, between different populations and their correlation with the variation of environmental features using an integrative approach combining metabolomics and geoclimatic data regarding the original location of each population (Meijón et al., 2016). This study used a collection of populations covering a significant portion of the species natural distribution, finding aridity to be the most determinant factor related to the natural variation observed at metabolic level, particularly that of secondary metabolism. Thus, this study defined two major population groups, Atlantic and Mediterranean, distinguished by secondary metabolism variation associated to the water availability they adapted to at their original geographical locations. However, further studies are necessary to improve our knowledge on the actual link between genetics, environmental conditions, and development and growth, as well as on the molecular processes behind the observed phenotypic natural variation.

In this context, integrative systems biology approaches are a good choice. They rely on the combination of different omic layers that complement each other providing a more comprehensive understanding of the studied biological process. They also allow to overcome limitations related to the lack of genetic or, in general, molecular information in the case of orphan species, as it is the case of maritime pine. This type of approach has been successfully used in the study of several processes related to pine trees biology, such as the response to UV (García-Campa et al., 2022; Jesus Pascual et al., 2017; J. Pascual et al., 2016), heat stress (Escandón et al., 2017, 2016; Lamelas et al., 2020; Roces et al., 2022), combined heat and drought stress (López-Hidalgo et al., 2023), needle development (Valledor et al., 2010), or apical bud maturation (Valledor et al., 2021).

In this manuscript we have generated proteomics data from three *P. pinaster* populations from the Northwest and Southeast of Spain, and Morocco, covering the main genetic pools of the Mediterranean basin, as described by Bucci et al., 2007) and showing phenotypical variation when grown in a common garden. The integration of this data with metabolomics, phenotypical and at-origin geoclimatic data on each population uncovered variation concerning several molecular mechanisms and their involvement with plant performance and development. Furthermore, two different organs, needles and apical and basal sections, were analyzed to cover a wider range of plant physiology and development, and to evaluate which one was more informative in terms of natural variation dynamics. Needle analysis showed the importance of photodamage in the Mediterranean populations and of ROS scavenging mechanisms and a metabolic rearrangement aimed at supporting secondary metabolism to deal with it and impacting growth. In turn, buds analysis provided insight on the importance of UV-B signaling and associated secondary metabolism for the protection of the shoot apical meristem (SAM) and of protein homeostasis and vesicle trafficking, potentially related to organ polarity, crucial in plant development, and therefore associated to tree architecture.

## Material and methods

### Plant Material and Growth Conditions

Plant material was sampled from three autochthonous populations of *P. pinaster* across the Mediterranean basin, corresponding to different climatic environments in relation to water availability, temperature, and irradiation (Figure 1). The populations selected were the following: Cadavedo (CDVD), Oria (ORIA) and Tamrabta (TAMR). CDVD is a population from the Cantabrian Sea area, in Northern Spain. This area is characterized by an Atlantic weather with an annual rainfall of 1316 mm quite uniformly distributed throughout the year. Consequently, the aridity index for CDVD provenance at its origin is 0.8. ORIA comes from Southern Spain, where annual precipitation is lower (357 mm) and the aridity index is 110. The altitude is also higher (1150 m), leading to a Direct Normal Irradiance (DNI) of 5935.384 Wh/m^2^/day. TAMR, a Moroccan provenance, comes from an area characterized by an annual precipitation of 763 mm and an aridity index of 67.9, both values in between those for CDVD and ORIA. TAMR location is also characterized by a high altitude (1760 m) and a DNI of 5565.384 Wh/m^2^/day (see Meijón et al., (2016) and Supplemental Table 1 for details on geoclimatic features corresponding to each provenance).

**Figure 1:**
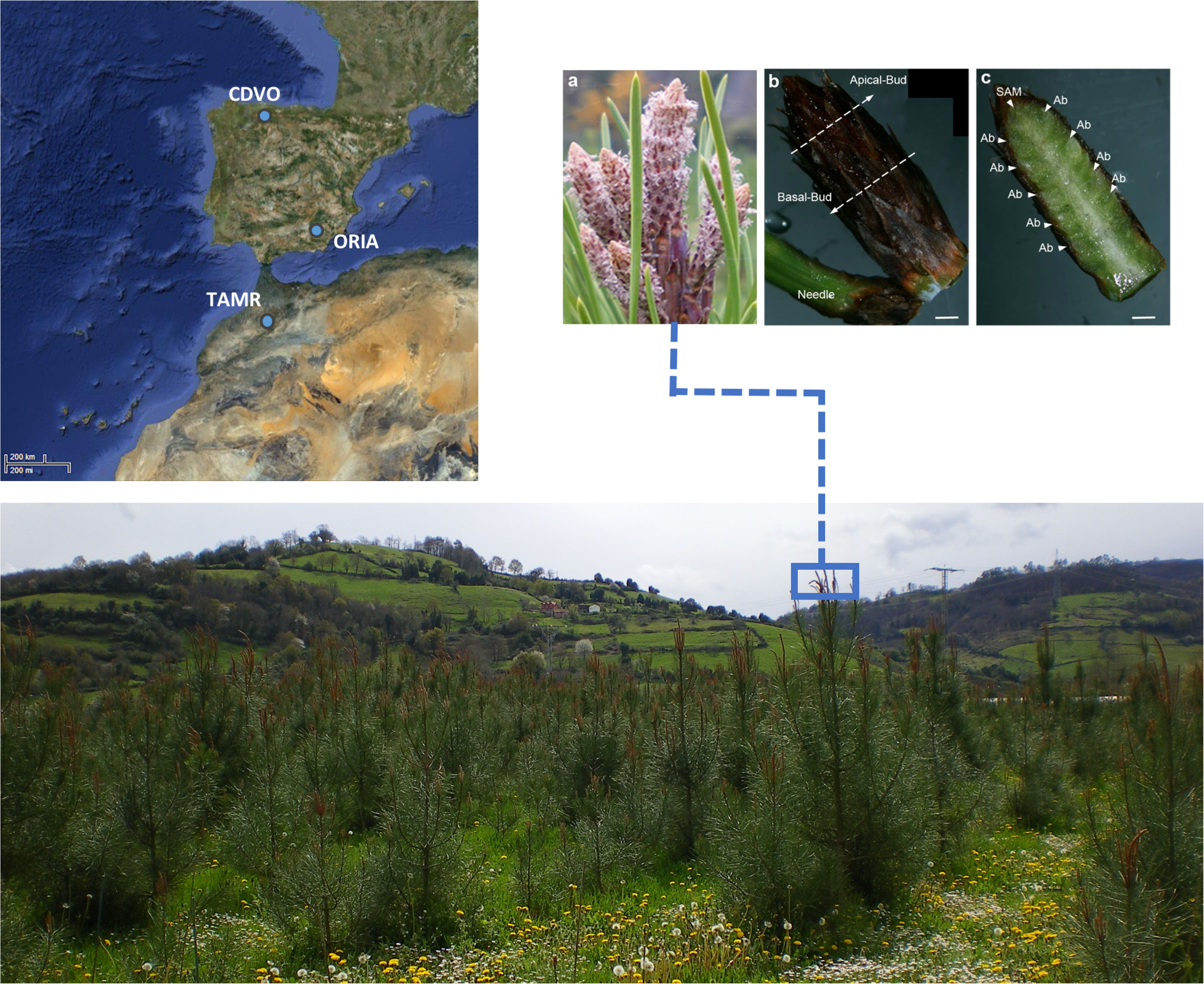
Provenances and organs/tissues overview. A. Natural localization of *Pinus pinaster* provenances used in the manuscript. B. Stage 1 of apical bud or budburst. C. Apical bud as seen under a stereoscope. D. Transversal section of an apical bud. SAM, shoot apical meristem; Ab, axillary bud. Scale bars = 1 mm.

For each provenance, seeds were collected from five mother trees separated from each other by at least 50 m to avoid inbreeding. Fifty open-pollinated families were available for the experiments. Seedlings were grown in a randomized block design in a common garden located in the Northwestern Spain (Experimental Station “La Mata”, Grado-Asturias: 43° 32′N 7°00′W, 65 m altitude range, 1100 mm annual rainfall, 12.5 °C annual temperature, 1.1 aridity index, 3032.654 DNI) Further details on this common garden design can be found in Gaspar et al., (2013). Plants were established in the field for 5 years and annual height (total growth), and number of whorls (polycylism frequency) were measured at the end of the summer corresponding to the end of the fifth year of growth (see Meijón et al., (2016) and Supplemental Table 1 for details on phenotypic features corresponding to each provenance). Sampling was done by triplicate as described in Meijón et al. (2016).

### Protein isolation, proteomic analysis and protein identification

Protein isolation was performed sequentially to the metabolite extraction performed in Meijón et al., 2016 using the multiple isolation protocol developed by Valledor et al., 2014.

Sixty μg of total protein were cleaned, digested, and desalted following the protocol described by Valledor and Weckwerth, 2014. Peptide chromatography and mass spectrometric analysis were performed according to Pascual et al., 2017a with a slight modification in the effective gradient: 90 min from 2 % to 50 % acetonitrile/0.1 % formic acid (v:v) with a column regeneration step of 22 min. A Chromoltih RP-18R 15 cm length 0.1 cm inner diameter column (Merck, Germany) was used.

Protein identification was performed with Proteome Discoverer 2.0 (Thermo Scientific, USA). Protein identification threshold was established at 5% and 1% false discovery rates (FDR) at peptide and protein levels, respectively. Only proteins with at least two identified peptides, one of them unique, were considered as identified. We used four databases for SEQUEST searches: *Pinus sylvestris* and *Pinus taeda* (34063 accessions) (Proost et al., 2014), and in-house built databases *Pinus pinaster* (117080 accessions) and *Pinus radiata* (67647 accessions), built following the procedure described by Romero-Rodríguez et al., 2014. Identified proteins were quantified by a label-free approach based on the estimation of the areas of the three most abundant peaks assigned to each protein by Proteome Discoverer. Proteins were functionally classified according to MapMan functional bins (Thimm et al., 2004) employing protein sequences and Mercator online tool v3.6 (Lohse et al., 2014). Protein annotation was performed using Sma3s (Casimiro-Soriguer et al., 2017).

Proteomics data are available via ProteomeXchange with identifier PXD048701.

### Metabolites re-identification

The metabolites obtained in Meijón et al., 2016 were re-identified (Supplemental Table 2 and 3). Only normalized peak areas (smoothed and deconvoluted) were maintained. The individual peaks were re-identified using Compound Discovered v.3.3.1 with a 3 ppm threshold and considering as “identified” those metabolites that were defined after the comparison to our standard compound library (containing >100 compounds) or by MS/MS matching to plant compounds for which their MS/MS is available in public databases (ChemSpider, Arita Lab 6549, EFS HRAM). Fragments present in collected MS/MS data were automatically matched with the predicted fragments generated using in silico fragmentation with a mass tolerance error of ± 3 ppm. Those compounds with FISh (Fragment Ion Search) coverages higher than 65% were included as “identified” (Jaén-Gil et al., 2018). Finally, molecular ions with exact masses corresponding to identified metabolites in public databases (KEGG, PubChem, METLIN, MassBank, HMDB and Plantcyc databases) were classified as “tentatively assigned” (with DeltaMass less than 5 ppm). Metabolite identification against our library was confirmed by retention time (RT), mass, isotopic pattern, and ring double bound parameters.

### Statistical and bioinformatics analysis

All statistical procedures were conducted with the R programming language running under the open source computer software R v4.3.0 (R Development Core Team, 2015) and RStudio v2023.03.1 (RStudio Team, 2016) using the packages pRocesomics^1^, MOFA2 (Argelaguet et al., 2020) and pheatmap (Kolde, 2019).

Proteomics and metabolomics data were pre-processed following the recommendations by Valledor and Jorrín (2011) and Valledor et al. (2014b). In brief, missed values were imputed using a Random Forest approach with an imputation threshold of 0.34, and variables were filtered out using a consistency filter of 0.34. Data were normalized by average intensity following a sample centric approach. Values were scaled and centered (z-scores) prior to univariate (one-way ANOVA followed by a Tukey HSD post-hoc test, P<0.05.) and multivariate analyses (PCA and sparse PLS) and heat map clustering. To avoid variable noise 15% variables with the highest variance coefficient were removed before MOFA. Cytoscape v. 3.7 (Shannon et al., 2003) was employed for network representation and analysis. Metabolites functional classifications was performed manually according to MapMan functional bins (Thimm et al., 2004).

## Results

### Needles and buds exhibited distinct proteomic profiles

Proteome natural variation was evaluated in three *P. pinaster* provenances with differential geographical origin, climatic conditions and growth characteristics grown in a common garden for 5 years (ORIA, CDVD and TAMR). In addition, we analyzed three different organs: needle, and bud apical and basal sections. Proteomic analysis allowed the identification of 2124 proteins, 2061 of which were suitable for quantification (Supplemental Table 4). A first overview of the proteins quantified in each organ clearly showed that the differences between them were rather quantitative, most of the proteins being coincident between them (Figure 2A). Principal component analysis (PCA) using the quantified proteins separated samples by tissue (Figure 2B; Supplemental Table 5), indicating major differences between samples are related to tissue type. A more detailed look into the top scoring proteins in principal component 1 (PC1), which separated samples by organ, showed chloroplastic proteins, especially photosynthesis-related proteins, among the proteins with the strongest positive correlation with needle samples (Supplemental Table 5). Regarding the proteins that showed the most negative correlation with PC1, where both apical and basal bud samples separated, included several proteasomal subunits, central metabolism enzymes and secondary metabolism enzymes like Cinammomyl-CoA reductase (Supplemental Table 5). Consistently, heatmap clustering of MapMan protein functional categories showed a clear higher abundance of photosynthesis proteins in addition to proteins related to RNA biosynthesis, including proteins belonging to the plastid and nuclear transcriptional machineries (Figure 2C). In turn, a cluster including polyamine and nucleotide metabolism, phytohormone action and protein modification was in general more abundant in bud samples in comparison with needles. Interestingly, the abundance of these categories was quite homogeneous in needle samples regardless of their provenance, but that was not the case in bud samples, showing more variant abundances between provenances as well as bud sections (Figure 2C). Moreover, 1669 proteins showed statistically differential abundance between organs (Supplemental Table 4).

**Figure 2:**
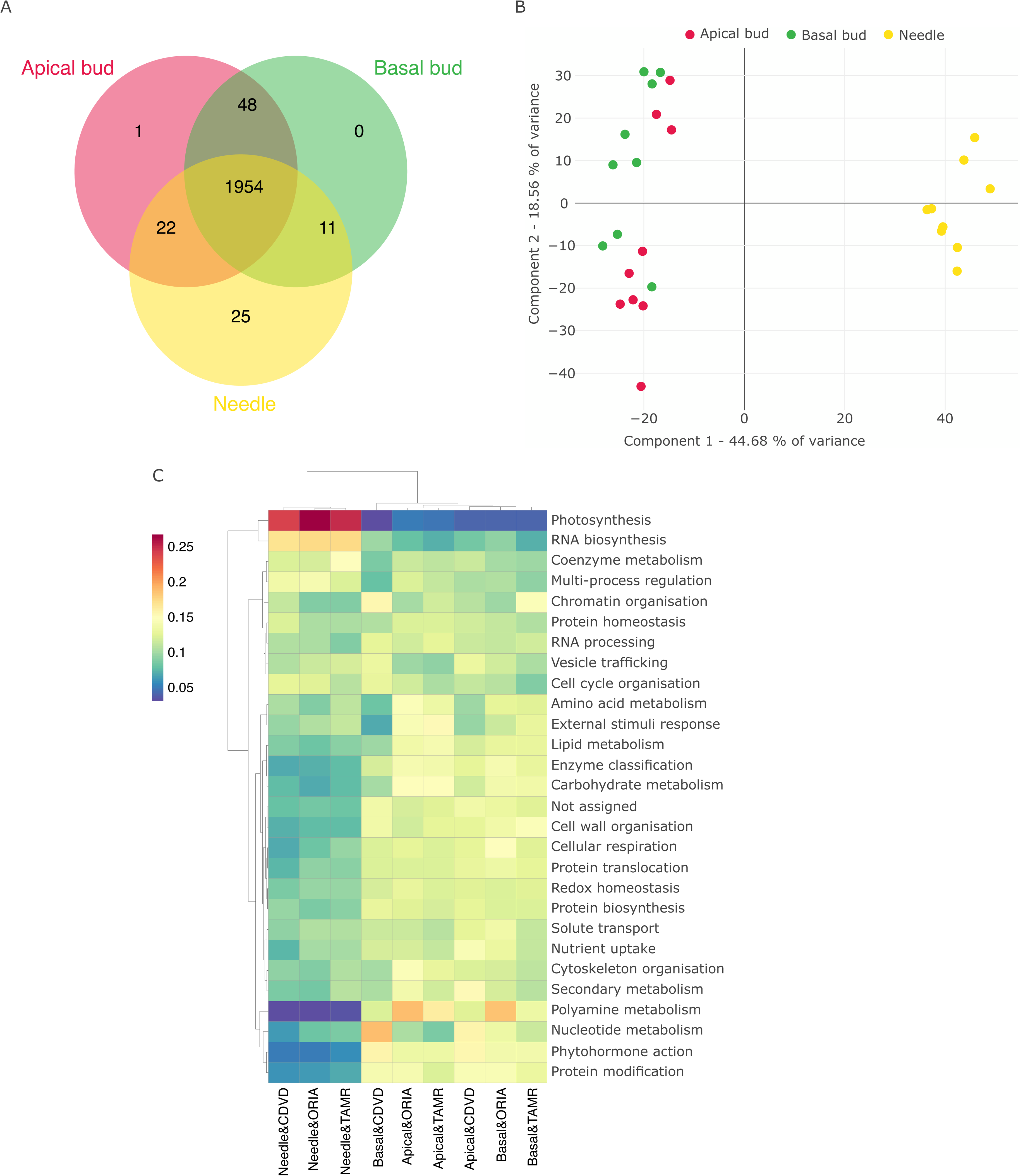
Proteomic analysis. A. Venn diagram showing the qualitative overlap between the proteins quantified in each tissue. B. Principal Component Analysis (PCA) plot showing first 2 principal components (PCs), explaining 63% of the observed variance. C. Heatmap clustering analysis of proteins according to their classification in MapMan categories. Distances were established employing Manhattan distance. Ward’s methods was used for aggregation. D. Heatmap clustering of protein z-scored quantitative values.

### Needle proteome characterization revealed provenance-specific functional signatures

As it was shown in previous section, the analysis of proteome natural variation was masked by major inherent differences in the proteins expressed in buds and needles. Therefore, we decided to analyze needles and buds proteomics data independently in order to explore the differences associated to the different provenances included in the study. Needle proteome dataset consisted of 1924 quantified proteins, 136 of which were differentially accumulated between provenances (ANOVA *P*-value <0.05; Supplemental Table 6). Venn analysis revealed minor qualitative differences between provenances. Most of the proteins were present in the three of them (Figure 3A; Supplemental Table 6). Therefore, the differences between provenances were mostly quantitative. Indeed, PCA of the 1924 quantified proteins showed a clear separation between the three provenances (Figure 3B; Supplemental Table 7). CDVD needles separated from ORIA and TAMR needles in PC1, while ORIA and TAMR needles separated along PC2 (Figure 3B). Heatmap clustering analysis of MapMan protein functional categories showed the most representative pathways of each provenance and the overall similarities between them (Figure 3C). Thus, CDVD needles clearly differed from ORIA and TAMR needles, which is in line with PCA results (Figure 3B). Regarding the abundance of specific functional categories, CDVD needles showed a higher abundance of proteins related to protein homeostasis and biosynthesis, chromatin organization, multi-process regulation, RNA processing and amino acid metabolism, and showed a particularly low abundance regarding cellular respiration, protein translocation, nutrient uptake and nucleotide metabolism (Figure 3C). TAMR and ORIA needles, on the contrary, showed in general a lower abundance of the categories that were more abundant in CDVD needles (Figure 3C). However, the accumulation of proteins from several categories, including photosynthesis, redox homeostasis, vesicle trafficking cellular respiration or nucleotide metabolism among others, was higher in TAMR and ORIA than in CDVD needles (Figure 3C). TAMR needles were characterized by a significant higher abundance of proteins related to amino acid metabolism, cytoskeleton organization, coenzyme metabolism, secondary metabolism, enzyme classification, protein modification, phytohormone action and external stimuli response (Figure 3C). These categories were, in turn, less abundant in CDVD and ORIA needles. In the latter, protein biosynthesis, carbohydrate and lipid metabolism, and RNA biosynthesis proteins were also less abundant than in needles from the other two provenances (Figure 3C).

**Figure 3:**
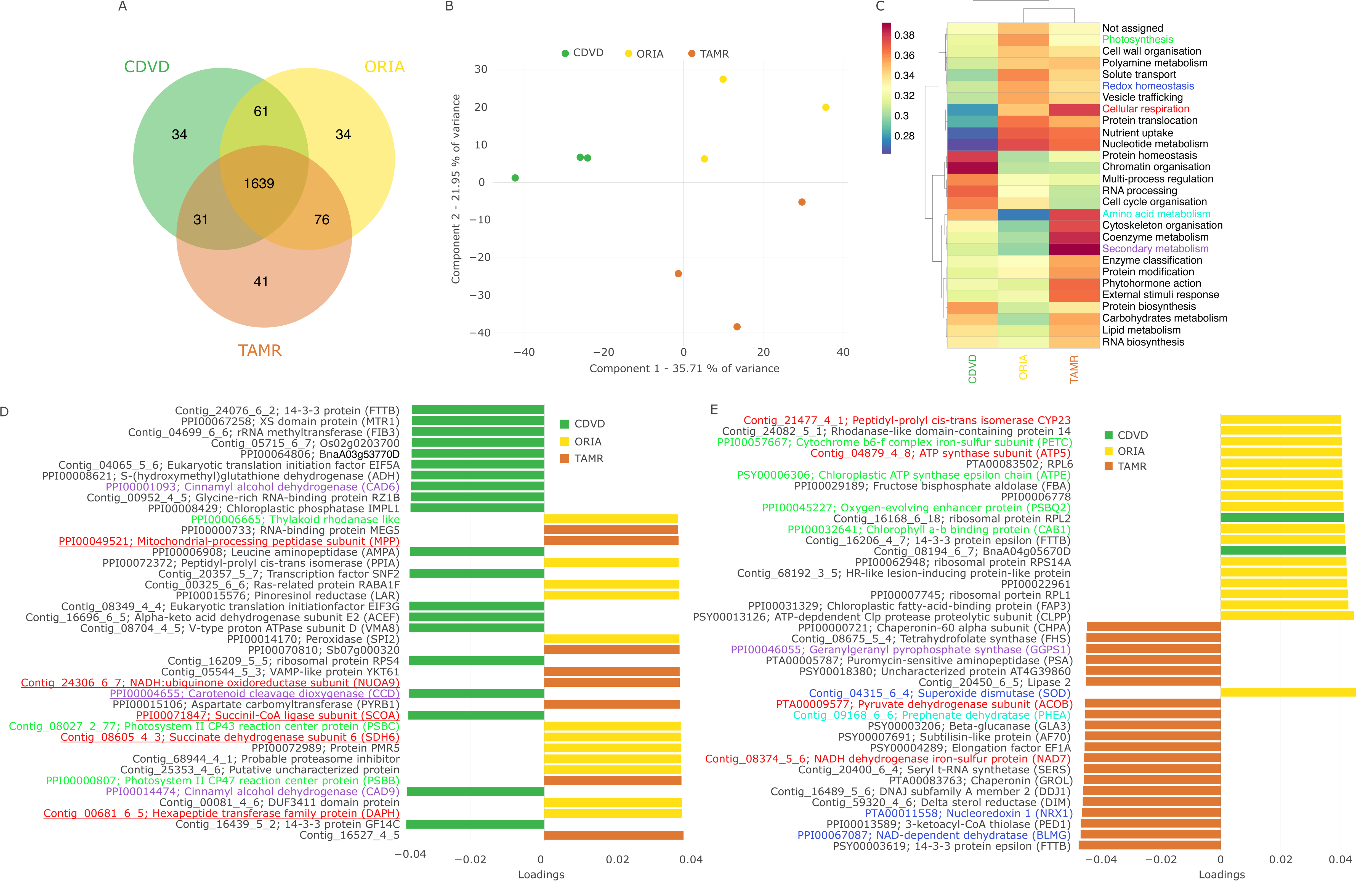
Needles proteomic analysis. A. Venn diagram showing qualitative comparison between the proteins quantified in needles from the three studied populations (CDVD, ORIA and TAMR). B. Principal Component Analysis (PCA) plot representing first two principal components (PCs), explaining more than 50% of the observed variance. C. Heatmap clustering of needle proteins MapMan functional categories abundance. Colored categories highlight those categories represented by several proteins among PCA top loading proteins, shown in D and E. D and E. PC1 and PC2 top loading proteins. Bar colors indicates the population that showed the highest abundance for each protein. Protein names colors refer to the MapMan functional categories highlighted in C.

The results of the analysis of MapMan functional categories were consistent with PCA top loading proteins (Figure 3D and E; Supplemental Table 7). According to differences in the abundance of photosynthesis-related proteins, PC1 top loading proteins included the photosystem proteins CP43 and CP47 (PSBC and PSBB), and a rhodanese-containing protein (STR4), required for anchoring ferredoxin-NADP reductase to the thylakoid membranes and sustaining linear electron flow (Figure 3D). These proteins showed the highest abundance in TAMR (Figure 3D). Also, in line with the MapMan analysis results, the cellular respiration proteins mitochondrial processing peptidase (MPP), subunits of the respiratory complex I (NUOA9) and II (SDH6), a TCA cycle succinyl-CoA ligase (SCOA), and an hexapeptide transferase (DAPH) were between the proteins with a higher loading (Figure 3D). With the exception of SCOA, they were more abundant either in ORIA or TAMR (Figure 3D). PC2 top loading proteins included further photosynthesis and cellular respiration proteins (Figure 3E). A cytochrome b6-f complex subunit (PETC), an oxygen-evolving enhancer protein (PSBQ2), an ATP synthase subunit (ATPE), and a chlorophyll-binding protein (CAB1) were more abundant in ORIA (Figure 3E). A pyruvate dehydrogenase subunit (ACOB) and an NADH dehydrogenase (NAD7), in turn, were more abundant in TAMR (Figure 3E). In addition to photosynthesis and cellular respiration, PC2 top loading proteins included proteins involved in redox homeostasis, and amino acid and secondary metabolisms (Figure 3E). A superoxide dismutase (SOD), involved in ROS scavenging, showed its highest abundance in ORIA, while a nucleorredoxin (NRX1) and an NAD-dependent dehydratase were more accumulated in TAMR (Figure 3E). TAMR was also the provenance showing the highest levels of a prephenate dehydratase (PHEA), part of the shikimate pathway, used for the biosynthesis of aromatic amino acids. Aromatic amino acids are the basis for plant secondary metabolism, also represented among PC2 top loading proteins by a geranylgeranyl pyrophosphate synthase (GGPS1), an enzyme involved in the synthesis of geranylgeranyl pyrophosphate, a precursor to carotenoids, tocopherols or gibberellins. GGPS1 showed the highest abundance in TAMR needles (Figure 3E).

### Bud proteomic analysis showed differences associated to bud section and provenance

The analysis of bud sections proteome quantified 2000 proteins, 968 out of which were differentially accumulated in the different samples (ANOVA *P*-value <0.05; Supplemental Table 8). Similarly to needle proteomes, we did not find major qualitative differences between samples regarding bud provenance or section (Figure 4A; Supplemental Table 8). PCA of the quantified proteins resulted in an effective separation of samples by provenance in PC2 and by section in PC1 (Figure 4B; Supplemental Table 9). Therefore, PC1 and PC2 represented predominantly the variance associated to provenance and bud section respectively. Noteworthy, as it happened in needle samples proteome PCA, CDVD samples grouped apart from the other two provenance in PC2 (Figure 4B).

**Figure 4:**
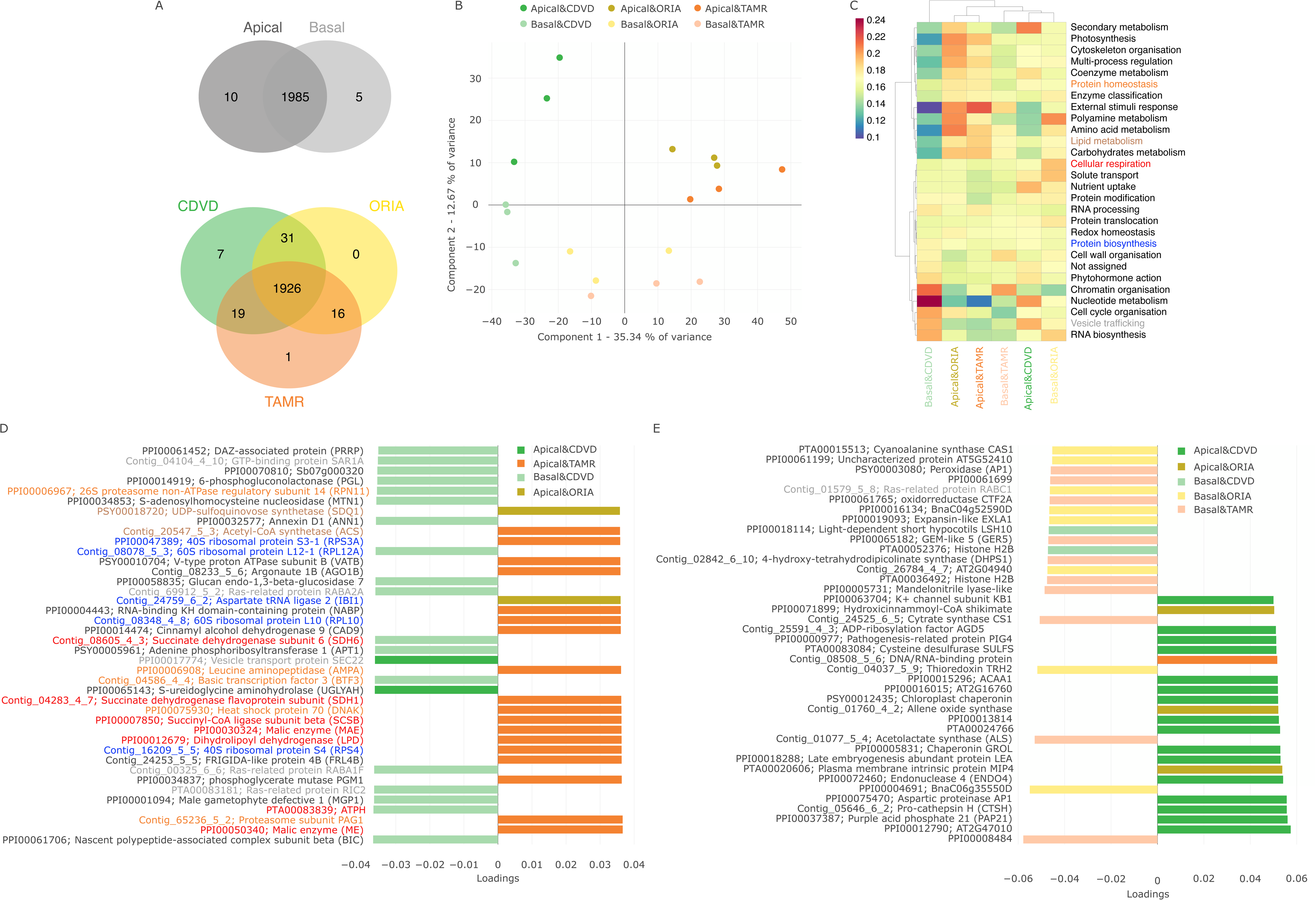
Buds proteomic analysis. A. Venn diagrams showing a qualitative comparison between the proteins quantified in buds by section and population. B. Principal Component Analysis (PCA) plot representing first two principal components (PCs), explaining 48% of the observed variance. C. Heatmap clustering of bud proteins MapMan functional categories abundance. Coloured categories highlight those categories represented by several proteins among PCA top loading proteins, shown in D and E. D and E. PC1 and PC2 top loading proteins. Bar colors indicates the population that showed the highest abundance for each protein. Protein names colors refer to the MapMan functional categories highlighted in C.

Analysis of the abundance of MapMan protein functional categories (Figure 4C) and PCA top scoring proteins (Figure 4D and E, Supplemental table 9) provided a broader overview on differential pathways between bud sections and provenances. PC1 top scoring proteins included several proteins related to cellular respiration, including tricarboxylic acid cycle proteins, and ribosomal and proteasome-associated proteins (Figure 4D). PC2 top scoring proteins, associated to the separation of samples by bud section, included a more heterogeneous group of proteins with a clear preponderance of proteins without functional annotation in MapMan ontology (Figure 4E). Regarding, MapMan functional bins, in general CDVD showed a lower abundance of proteins related to external stimuli, polyamine, amino acid and carbohydrate metabolisms (Figure 4C). CDVD basal buds was the most different sample, characterized by a higher abundance of proteins related to chromatin organization or RNA biosynthesis, and a significant lower abundance of several functional categories, including secondary metabolism, photosynthesis and lipid metabolism in addition to the above-mentioned categories, common to both CDVD bud sections (Figure 4C).

In concordance with these results, PC1 top scoring proteins included an Argonaute protein (AGO1B), involved in post-transcriptional silencing and ncRNA-mediated epigenetic regulation, and S-adenosylhomocysteine nucleosidase (MTN1), part of the activated methyl cycle and therefore an integral component in methionine metabolism and regulator of the trans-methylation capacity, relevant for epigenetic regulation by DNA methylation (Figure 4D). Similarly, histone H2A was among PC2 top scoring proteins (Figure 4E). The pathway abundances observed in CDVD basal buds were in general the opposite in TAMR and ORIA basal buds, which interestingly clustered along with CDVD apical buds (Figure 4C). ORIA and TAMR apical buds showed a rather similar pathway abundance pattern with a more accused contrast with CDVD basal buds (Figure 4C).

### Integrative proteomics and metabolomic analysis identified the main functional sources of natural variation

To advance our research and attain a more thorough understanding of the molecular processes associated to *P. pinaster* natural variation, we integrated pre-existing metabolomics data from the three studied provenances (Meijón et al., 2016; Supplemental Table 2 and 3) with the proteomics data generated for this study. Employing Multiomic Factor Analysis (MOFA), we were able to delve deeper into the significance of the proteomic and metabolomic layers and the interplay between them in the context of the natural variation captured at each level. Following the same approach as in the analysis of the proteomics data, different tissues were analyzed separately.

Needles data MOFA inferred 5 latent factors (LFs) representing sources of variation across the data (Figure 5A). Overall, these LFs represented 85% of proteome total variance and 61% of metabolome variance (Figure 5A; Supplemental Table 10). Nonetheless, each LF showed a different contribution of each molecular level (Figure 5A; Supplemental Table 10). A more detailed analysis of each factor showed that LF1 and LF2 allowed discriminating samples by their provenance (Figure 5B and C). These LFs were further characterized.

**Figure 5:**
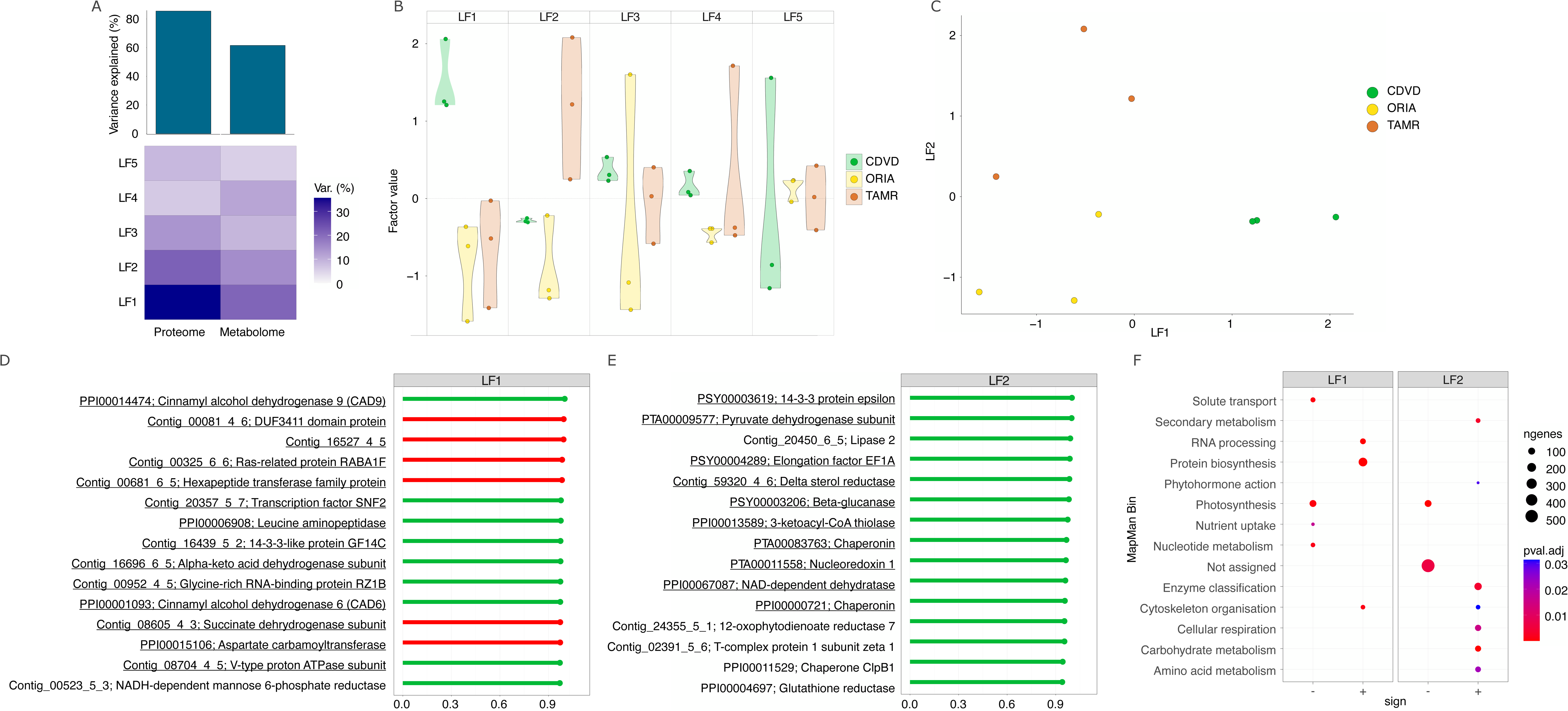
Needles Multi-Omics Factor Analysis (MOFA). A. MOFA explained variance plot depicting the variance explained at proteome and metabolome levels and its distribution through the five latent factors (LFs) inferred by the analysis. B. Visualization of needle samples distribution in each inferred LF. C. Visualization of needle samples distribution by the combination of LF1 and LF2. D and E. MOFA LF1 and LF2 top weight features. Green and red bars represent positive and negative weights respectively. Underlined features highlight proteins that were between needles PCA top loading proteins (Figure 2D and E). F. Gene set enrichment analysis of LF1 and LF2 associated features using MapMan ontology. Dot sizes represent the number of genes. Dot colors represent the adjusted p-value calculated by the performed gene set enrichment analysis.

Top weight features in both LF1 and LF2 were proteins (Figure 5D; Supplemental Table 10) consistently with the higher contribution of the proteome to the explained variance captured by each LF: a 35% and a 21% versus a 20% and a 14% in the case of the metabolome (Figure 5A; Supplemental Table 10). LF1 top weight features included 13 proteins that were between the PC1 top loading proteins (Figure 5D; underlined), including several related to cellular respiration, identified as a main source of variation between provenances in the performed PCA (Figure 3). Overall, this overlapping is in concordance with the separation of CDVD samples by both PC1 and LF1. The overlapping between LF2 and PC2 was similar in number: 11 out of the 15 LF2 top weight proteins were among PC2 top loading proteins (Figure 5E; underlined). These proteins included a pyruvate dehydrogenase subunit, involved in cellular respiration, and the redox homeostasis proteins NAD-dependent dehydratase and nucleoredoxin 1, all of them top loading proteins in PC2 as well (Figure 3E). In addition, MOFA LF2 top weight features included a glutathione reductase, also involved in redox balance. The signs of MOFA weights and PCA loadings were also in concordance.

Next, we performed a functional enrichment analysis following Mercator 4 functional annotation to further investigate the pathways represented by the features included in LF1 and 2 (Supplemental Table 11). This analysis revealed a statistically significant representation of features related to RNA processing, protein biosynthesis and cytoskeleton organization among the features with a positive weight (Figure 5F). In turn, LF1 features negative counterparts were enriched in photosynthesis, nucleotide metabolism, solute transport and nutrient uptake categories (Figure 5F). Features with a positive weight in LF2 were enriched in cellular respiration, carbohydrates and amino acid metabolism, secondary metabolism, phytohormone action and enzyme classification (Figure 5F). Negative-weight LF2 features were enriched in photosynthesis-related and not assigned features (Figure 5F). Altogether, MOFA results strength PCA results, further supporting cellular respiration, photosynthesis and secondary metabolism as relevant sources of natural variation between the three provenances.

Following the same approach with bud samples, MOFA inferred 4 LFs (Figure 6A). As it happened in the analysis of needle samples, the proteome represented an overall higher variance than the metabolome, although the percentage of each omic layer varied between LFs (Figure 6A; Supplemental Table 10). In the case of buds, LF1 accomplished to capture the differences between CDVD apical and basal buds (Figure 6B and C), emphasizing again the singularity of CDVD buds, as it was previously evidenced by the proteomic analysis. In concordance with the higher variance explained by the proteome in LF1, LF1 top weight features were all proteins (Figure 6D). Most of them were among buds PCA PC1 top loading proteins (Figure 6D; underlined). LF2 contained the variance associated to bud section (Figure 6B and C), which was also captured in proteomics data PCA (Figure 4B). In this case, top weight proteins included mostly metabolites as the variance explained by the metabolome in LF2 was higher than that of the proteome (Figure 6E). It also suggests that main differences between apical and basal buds are reflected at the metabolome rather than at proteome level. LF2 top weight metabolites are mostly unidentified, with the exception of propylmalic acid, pipericine and dihydroxychalcone, all of them secondary metabolites and more abundant in apical buds according to their positive weight in LF2 (Figure 6E).

**Figure 6:**
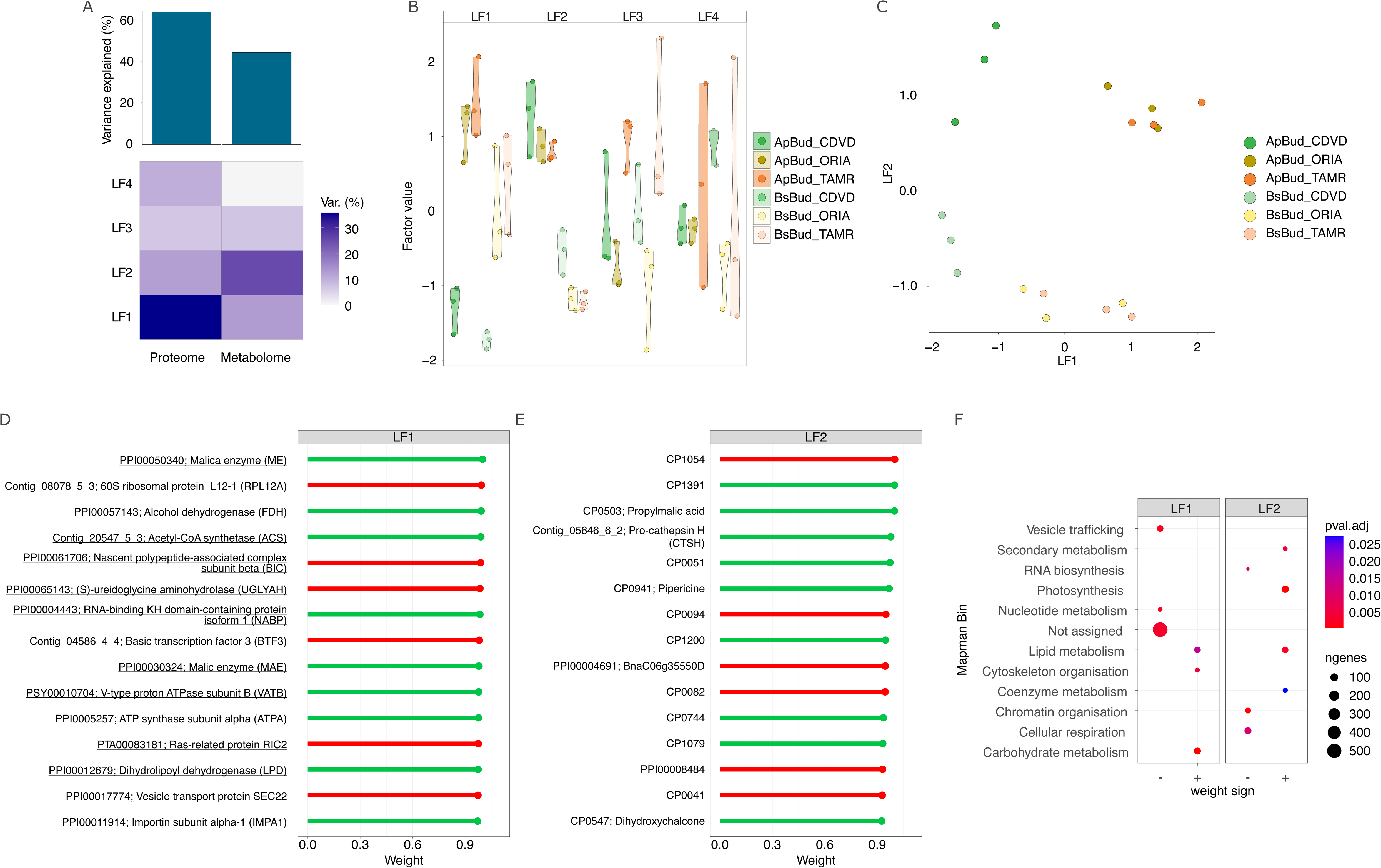
Buds Multi-Omics Factor Analysis (MOFA). A. MOFA explained variance plot depicting the variance explained at proteome and metabolome levels and its distribution through the four latent factors (LFs) inferred by the analysis. B. Visualization of bud samples distribution in each LF. C. Visualization of bud samples distribution by the combination of LF1 and LF2. D and E. MOFA LF1 and LF2 top weight features. Green and red bars represent positive and negative weights respectively. Underlined features highlight proteins that were between buds PCA top loading proteins (Figure 3D and E). F. Gene set enrichment analysis of LF1 and LF2 associated features using MapMan ontology. Dot sizes represent the number of features. Dot colors represent the adjusted p-value calculated by the performed gene set enrichment analysis.

An enrichment analysis of MOFA features, both proteins and metabolites, associated to LFs was performed to get a wider functional perspective of the processes and pathways most significantly represented in each case (Supplemental Table 11). LF1 features with a positive weight were significantly enriched in vesicle trafficking, nucleotide metabolism and proteins not assigned to any MapMan bin (Figure 6F). The negative ones, correlated with CDVD samples, showed an enrichment in lipid and carbohydrates metabolism, and cytoskeleton organization (Figure 6F). These results are congruent with the proteome PCA and MapMan abundances analysis (Figure 4). Regarding LF2, representing the variance between bud sections, negative features were enriched in RNA biosynthesis, chromatin organization and cellular respiration, while the positive ones were enriched in features related to secondary metabolism, photosynthesis, lipid and coenzyme metabolism (Figure 6F). The importance of variation concerning secondary metabolism was evidenced by buds proteomics PCA and MapMan analysis (Figure 4) as well as the presence of several identified secondary metabolites between LF2 top weight features (Figure 6E).

### Needles sPLS network revealed natural variation associated to secondary metabolism

MOFA allowed to discriminate between organs and provenances as well as to identify the most differential pathways according to the integration of proteomics and metabolomics data. A wider system-like view aimed at better understanding the correlations between individual proteins and metabolites and with environmental parameters of the different provenances in their original geographical locations was accomplished by sPLS network analysis. We used proteomics data as predictor for metabolomics and environmental data related to the origin of the provenances to gain detailed insight into the correlations between these features, especially those concerning the environmental origin differences between the analyzed provenances, and the pathways they belong to.

The high-confidence network obtained using needles data had several clusters of proteins centered on one or more environmental factors and/or metabolites, especially secondary metabolites (Figure 7). The accumulation of the proteins and metabolites in these clusters differed between provenances (Figure 7).

**Figure 7:**
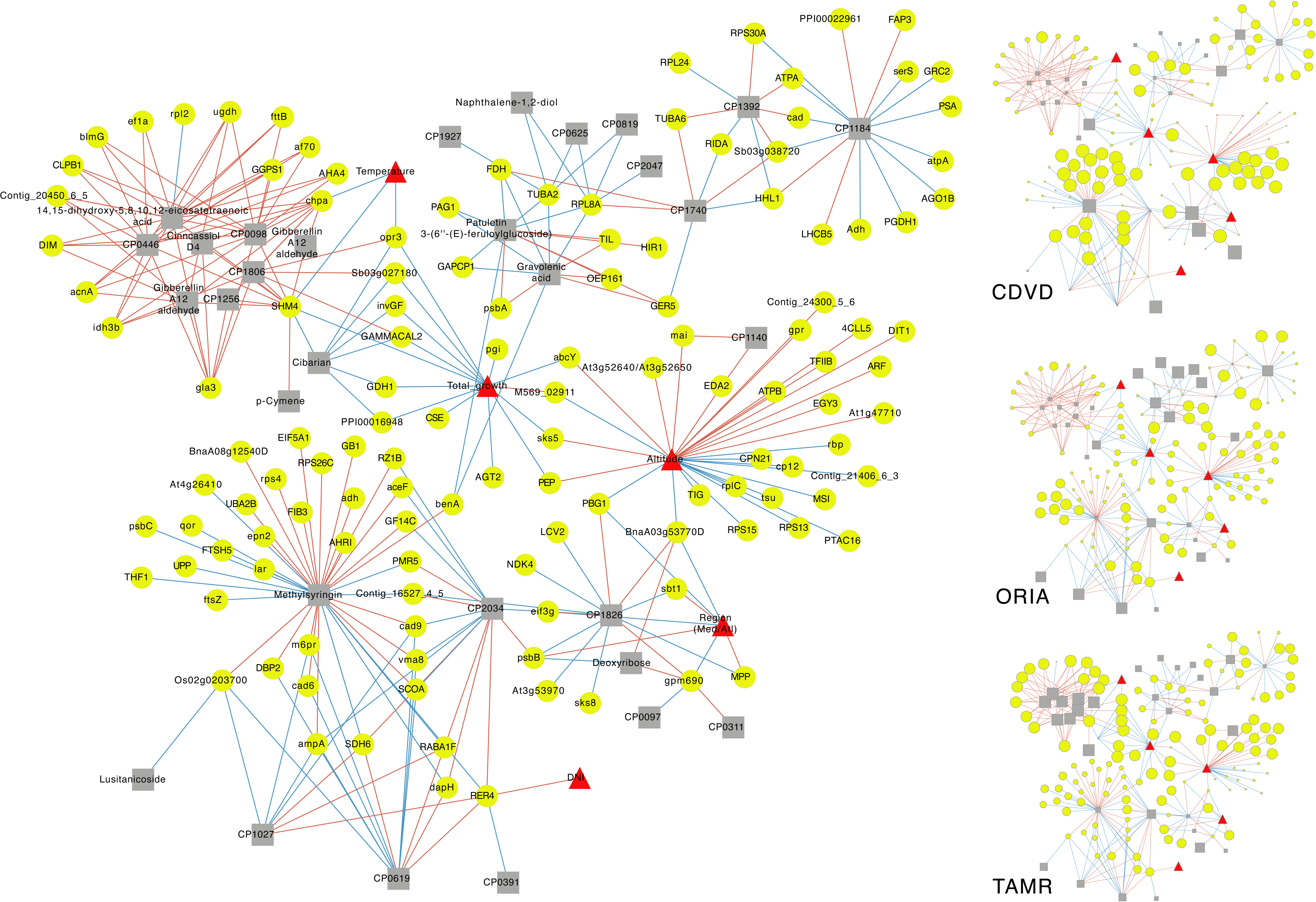
Needles integrative analysis. Needles sPLS-based network built using proteins as predictor for metabolites, and phenotypic and geoclimatic data. Correlation cutoff was set to 0.9. Red and blue edges represent positive and negative correlations respectively. Edge thickness increases with correlation absolute value. Proteins are represented by yellow circles, metabolites by gray squares and phenotypic and geoclimatic features by red triangles. Smaller networks beside the main one represent z-scored abundances of each feature in each population as node size.

A main cluster was organized around several metabolites, including gibberellins, cinnacassiol, eicosatetranoid acid and several unidentified metabolites. These metabolites had a strong positive correlation with proteins involved in redox homeostasis, protein biosynthesis, lipid metabolism and cellular respiration, which is consistent with the higher abundance of these functional categories in TAMR proteome (Figure 3C). Cellular respiration was also represented by several proteins in MOFA LF2 top positive weight features, which separated TAMR from ORIA and CVDO (Figure 5), as well as in PCA top loading proteins (Figure 3D and E). In concordance with the positive correlation between the proteins and the metabolites in the cluster, the higher abundance of the metabolites was found in TAMR, where the levels of the correlated proteins were higher than in the other two provenances, which showed lower levels of both the proteins and metabolites in the cluster (Figure 7). Regarding gibberellins, PC2 top loading proteins included GGPS1, responsible for the synthesis of geranygeranyl pyrophosphate, used as gibberellin precursor and that showed a higher abundance in TAMR (Figure 3E).

Some of the proteins in the previous cluster correlated inversely to the average temperature at the places of origin of the three provenances (CDVD, 13.2 °C; ORIA, 13 °C; TAMR, 10.8 °C): the chaperonin subunit CHPA, involved in protein folding; the methyl transferase SHM4, involved in several processes, like photorespiration; and an oxophytodienoate reductase (OPR1/3), involved in oxylipin and jasmonic acid metabolism. OPR1/3 showed also a negative correlation with an unidentified metabolite and total growth, showing the highest abundance in TAMR, which is the provenance with the lowest total growth (36.4 m vs 44.6 m of ORIA and 48 m of CDVD). Total growth was inversely correlated with other proteins, all of them again more abundant in TAMR, and including some metabolic enzymes, like an invertase (INVGF), a carbonic anhydrase (GAMMACAL2), a glucose 6-phosphate isomerase (PGI) and several amino acid metabolism enzymes: a glutamate dehydrogenase (GDH1), that generates 2-oxoglutarate (2-OG), an alanine-glyoxylate aminotransferase (AGT2) and an adenosylhomocysteine hydrolase (ABCY). Differences regarding amino acid metabolism had been already revealed by MapMan analysis (Figure 3C) and PCA (Figure 3D and E). Consistently with the inverse correlation between the abundance of these enzymes and total growth, the abundance of the mentioned enzymes was found to be the lowest in CDVD, which is the provenances with the largest total growth. This cluster included also a couple of proteins related to the cell wall: a caffeoylshikimate esterase (CSE), an enzyme in the lignin biosynthetic pathway (Vanholme et al., 2013), and SKS5, an extracellular glycosyl phosphatidylinositol-anchored glycoprotein (Sedbrook et al., 2002). Some of these proteins correlated inversely with total growth, but directly with the altitude at the locations of origin of the three analyzed provenances. Thus, they were less abundant in CDVD, naturally growing at 180 m over sea level, in comparison with ORIA or TAMR, naturally growing at altitudes over 1000 m. Also directly correlated to altitude, we found the chloroplastic transporter DIT1, a 2-OG/malate antiporter, and proteins related to phenylalanine metabolism: a maleylacetoacetate isomerase (MAI), involved in Phe degradation into fumarate and a 4-coumarate-CoA ligase (4CLL5), part of the phenylpropanoids pathway, which generates precursors for a wide range of secondary metabolites, including flavonoids, anthocyanins, lignin and sinapate esters among many others. Related to sinapate esters, a serine carboxypeptidase (EDA2) was included in this cluster postitively correlating with altitude. This is consistent with the role of serine carboxipeptidases in the biosynthesis of sinapate esters associated with the protection against UV radiation, increasing with altitude and a main cause of oxidative stress at several chloroplasts level. The presence of the chloroplast metalloprotease ethylene-dependent gravitropism-deficient and yellow-green EGY3, known as a mediator of chloroplast ROS homeostasis and retrograde signalling, positively correlating with altitude suggested increased levels of oxidative stress with altitude. The abundance of an ATPase subunit (ATPB) also correlated with altitude and varied along with the mentioned amino acid and secondary metabolism enzymes, suggesting changes in energy metabolism associated with modulations in secondary metabolism with altitude, what supports the relevance of secondary metabolites and highlights them as a main source of natural variation between the needles of the studied *P. pinaster* provenances.

The importance of secondary metabolites is also supported by their presence in the high-confidence sPLS network. Patuletin, an O-methylated flavonol, and gravolenic acid, an hydroxycinnamic acid, i.e., a phenylpropanoid, formed a cluster. Both metabolites showed a higher abundance in ORIA (Figure 7). In concordance with their photoprotective role, they correlated positively with the photosystem II reaction center protein PSBA and the outer envelope pore protein OEP161, responsible for importing PROTOCHLOROPHYLLIDE OXIDOREDUCTASE A (PORA) into chloroplast and thereby related with photoprotection (Samol et al., 2011). In addition, both metabolites had a positive correlation with plant defense and immunity proteins (subtilisin TIL, hypersensitive response protein HIR1, germin-like protein GEM5). Another phenylpropanoid included in the network showing an important connectivity with a good number of proteins was methylsyringin, derived from syringin, also called sinapyl-4-O-glycoside and related to plant immunity (Hemm et al., 2004). Syringin level has been also reported to increase under drought in several species (Liang et al., 2021). Therefore, the computed network pinpointed correlations between primary and secondary metabolism and contextualize them within differences in environmental factors at the original geographical locations of the studied provenances. Furthermore, it also hinted their correlation with stress factors, suggesting potential differences in stress response between provenances.

### Two sPLS networks deepen bud section and provenance natural variation

The high-confidence sPLS network generated using buds data resulted in 2 subnetworks (Figure 8). One of them included features which abundance varied mostly between bud section (Figure 8A), while the variation in the abundance of the features in the second one was mainly due to provenance (Figure 8B). In line with it, most of the metabolites in the first network were MOFA LF2 top weight features, related to variance between apical and basal bud sections (Figure 6). Only one metabolite in this network was identified: propylmalic acid, a terpenoid with an osmoregulatory role under drought (Liang et al., 2021). Propylmalic acid was unraveled as a top weight metabolite in MOFA LF2, which captured the variance between apical and basal bud sections (Figure 6). This compound was more abundant in apical buds in general and in those from TAMR in particular, thus showing natural variation associated to plant provenance as well (Figure 8A). Propylmalic acid derives from malic acid, synthesized by malic enzyme. Two of them were among the top loading proteins in buds PCA PC1 (Figure 4D) and top weight MOFA LF1 (Figure 6D). They were also more abundant in TAMR (Figures 4D and 6D). The abundance of the metabolites in this subnetwork was, whit the exception of CP1736, CP1054 and CP0737, higher in apical than in basal buds and correlated directly with the accumulation of several proteins related to the response to environmental and stress factors, like molecular chaperons (GROL and HSP2) or an allene oxide synthase (AOS), related to oxylipins synthesis. AOS was a PCA PC2 top loading protein as well and its abundance was higher in ORIA apical buds (Figure 4E). Oxylipins derive from fatty acids, synthesized in chloroplasts from acetyl-CoA. In this regard, buds PCA PC2 top loading proteins included an acetyl-CoA synthetase (ACS) and a UDP-sulfoquinovose synthetase (SDQ1), both involved in lipid metabolism and more abundant in ORIA apical buds (Figure 4D). Altogether, these results suggest natural variation associated to lipid metabolism associated to oxylipins. In addition, this cluster also included the UV signaling protein UVR8, that showed a higher abundance in ORIA apical needles. UVR8 is involved in UV-B signaling. It is a main component in UV-B-induced photomorphogenic responses that determine plant architecture (Huang et al., 2014; Jenkins, 2014).

**Figure 8:**
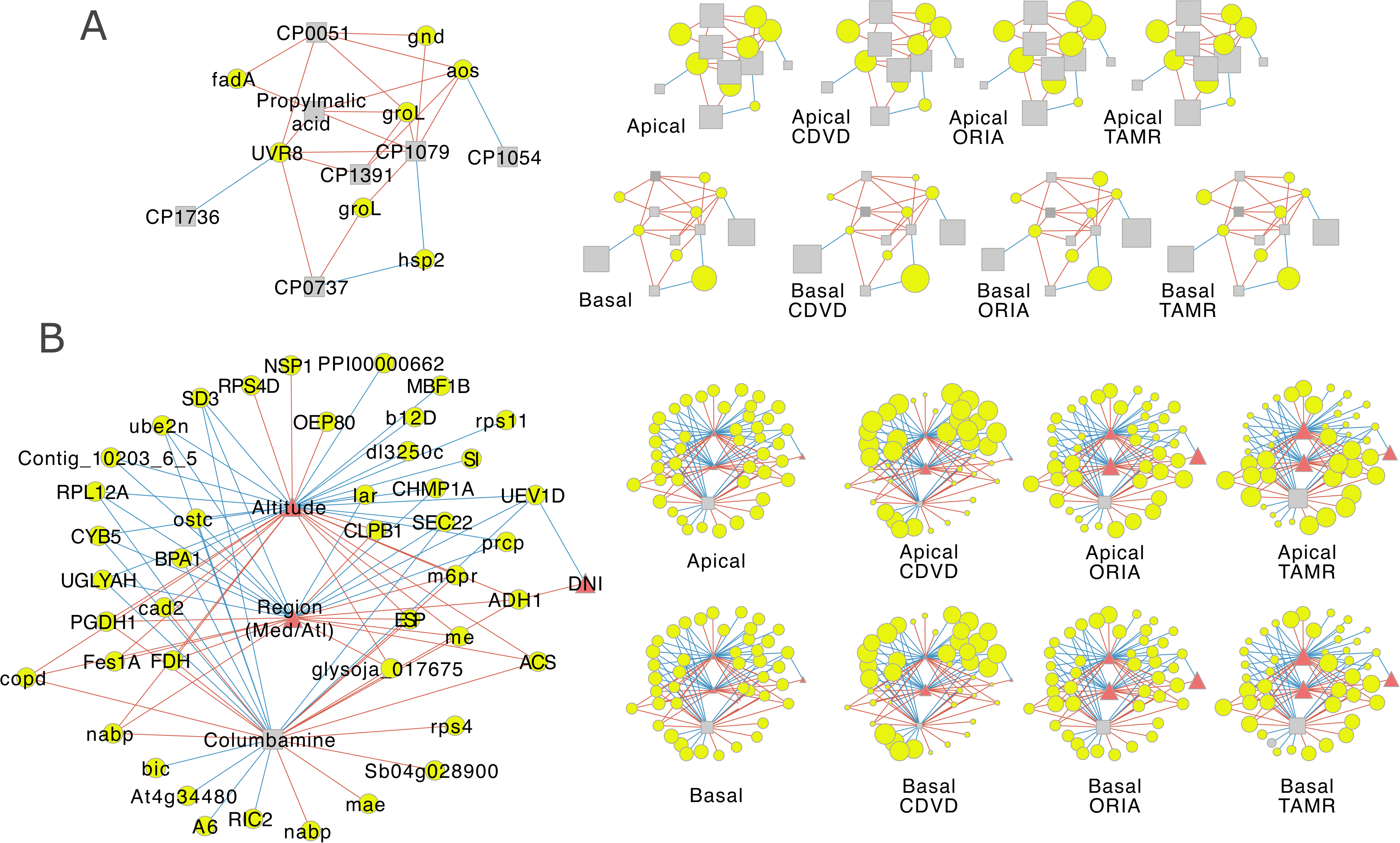
Buds integrative analysis. A and B. Buds sPLS-based networks built using proteins as predictor for metabolites, and phenotypic and geoclimatic data. Correlation cutoff was set to 0.85. Red and blue edges represent positive and negative correlations respectively. Edge thickness increases with correlation absolute value. Proteins are represented by yellow circles, metabolites by gray squares and phenotypic and geoclimatic features by red triangles. In the smaller networks besides each main network, node sizes represent z-scored abundances of each feature by bud section or by bud section and provenance.

The second high-confidence sPLS network was organized around Altitude and Region (Med/Atl) as environmental factors, and the metabolite columbamine, an alkaloid (Figure 8B). As already mentioned, this network contained mainly variation between provenances rather than between bud sections. However, it included proteins like ACS and a malic enzyme (ME), previously mention in the context of the other network and top proteins in both PCA and MOFA (Figures 4 and 6). These proteins had a direct correlation with the three features (Figure 8B). The abundance of a second malic enzyme (MAE), also unraveled as relevant by PCA and MOFA (Figure 4 and 6), correlated positively with the abundance of columbamine (Figure 8B), a secondary metabolite increasing under UV-B exposure (Liu et al., 2022).

## DISCUSSION

Natural variation within a species is a consequence of genetic differences derived from prevailing natural selection pressure in different geographical locations/ecological niches with different environmental or geoclimatic features. Studying natural variation requires growing different provenances of a species in a common garden under the same environmental conditions. This way environmental effect is minimized or standardized, and differences due to genetic effects can be better determined. In this study, we have analyzed three maritime pine populations. The selected provenances are classified in Atlantic (CDVD) and Mediterranean (ORIA and TAMR), regions differing in their average annual rainfall and aridity index. The three locations differ also in their altitude and Direct Normal Irradiance (DNI), increasing with altitude. The adaptation of each population to the geoclimatic conditions at their original locations gives different tree architectures as a result (Leyre Corcuera et al., 2012). These are genetically determined to a wide extent as they are observed when the three provenances were grown in a common garden, under the same conditions. Provenances showed different degrees of polycyclism as well, what determines branching, number of whorls and height, and therefore the architecture of the tree trunk (Supplemental Table 1). Everything considered, CDVD trees grow taller than TAMR trees. However, TAMR tree trunk is more valued by the timber industry for their lower number of nodes. ORIA trees are in a middle situation regarding CDVD and TAMR individuals. Similar trends have been observed under greenhouse conditions (Sánchez-Gómez et al., 2010).

Regarded the phenotypical differences between the three studied provenances, observed in a common garden, in which the conditions were the same for all of them, we aimed to decipher the molecular basis of the observed natural variation using proteomics and integrating it with available metabolomics, quantitative geoclimatic features at provenances original locations and quantitative growth and development parameters corresponding to the three provenances. Furthermore, the analysis of needles and two bud sections, basal and apical, allowed capturing molecular information on general plant performance (needles), related to growth and stress response, and plant development (buds), related to tree architecture.

The employment of a proteomic approach allowed to classify the samples by organ in a PCA (Figure 2A). However, it did not accomplish a full separation of bud samples by section, unlike the metabolomic analysis performed in a previous publication (Meijón et al., 2016). In fact, metabolic changes presented a rather qualitative nature, existing a high number of tissue-specific metabolites. Our proteomic analysis showed quantitative differences instead, i.e. the set of proteins expressed in each tissue did not differ drastically, but they showed different abundances (Figure 2B, C and D). Needles are, unlike buds, a photosynthetic organ. This was evidenced in the performed proteomic analysis when analyzing the abundance of MapMan functional categories (Figure 2D). Therefore, the larger source of variation captured by the proteome corresponded to the different nature of the analyzed organs. Gaining resolution on differences associated to provenance required analyzing organs independently. This strategy allowed a successful classification of needles by provenance (Figure 3B) and buds by both provenance and section (Figure 4B).

Needles, as already mentioned, are photosynthetic organs, providing energy to sustain the metabolism of the entire plant. The maintenance of adequate photosynthesis rates is crucial for plant physiology and largely determines plant growth (Long et al., 2006; Keller et al., 2022). Photosynthetic machinery is very sensitive to environmental conditions, including light intensity and spectral quality or water availability (Ashraf and Harris, 2013; Chan et al., 2010; Mayank Anand Gururani et al., 2015). It is susceptible to protein damage leading to decreased photosynthetic rates and energy imbalances producing oxidative stress and reducing growth (Li and Kim, 2022). Dealing with these phenomena requires photoprotection and ROS scavenging mechanisms (Foyer, 2018). Our proteomic analysis revealed differences at photosynthesis and redox homeostasis level as an important source of variation between provenances that is in concordance with the harshness of the environmental conditions they evolved in and are adapted to. Proteins belonging to these functional categories were more abundant in ORIA and TAMR (Figure 3C), both from areas with a high incidence of drought periods and extreme temperatures. Their importance was highlighted by the presence of several proteins from these categories in the list of PCA top loading proteins (Figure 3D and E) and MOFA top weight features (Figure 5D and E), which have a higher mathematical importance in the separation of samples by provenance in these analyses. The abundance of photosynthesis proteins is important for photosystem biogenesis, assembly and repair, especially in the case of PSII (Li et al., 2018, 2020), including proteins like PSBC, PSBB or PSBQ, all of them found to be more abundant in ORIA and TAMR (Figure 3D and E). Intraspecific variation in PSII photochemical efficiency has been previously reported in maritime pine in response to low temperatures (Corcuera et al., 2011). Photosystem proteins are especially prone to damage under stress conditions, including drought stress (Mayank Anand Gururani et al., 2015), associated to low annual rainfall and high aridity index, which would be in concordance with the higher abundance of these proteins in ORIA and TAMR. The abundance of these proteins could be also a consequence of the higher DNI ORIA and TAMR are adapted to. The acclimation to higher irradiances courses with increases in the content of photosynthesis proteins (Schöttler and Tóth, 2014). These could be an alternative explanation for the higher abundance of photosynthesis proteins in ORIA and TAMR, although it does not exclude higher reparation rates due to photodamage. Regardless of the cause, oxidative stress is consequently to photodamage (Foyer, 2018). Transient increases of ROS are a central phenomenon in retrograde signaling (Chan et al., 2016), but prolonged high ROS levels are detrimental for plant growth and development (Mhamdi and Van Breusegem, 2018). ROS levels are kept at bay by ROS scavenging machinery. In concordance with a possible increased photodamage, redox homeostasis proteins were more abundant in TAMR and especially in ORIA, coincident in this case with the higher abundance of photosynthesis proteins in this provenance (Figure 3C). In fact, superoxide dismutase, a main ROS scavenging enzyme, catalyzing the conversion of superoxide into hydrogen peroxide, was more abundant precisely in ORIA and was a top loading protein in the PCA (Figure 4E). This would be consistent with a presumable higher incidence of photodamage in ORIA as thylakoid membrane is a major site of superoxide production (Foyer and Hanke, 2022).

In addition to an increase abundance of ROS detoxification enzymes, TAMR showed an increased abundance in proteins related to secondary metabolism (Figure 4C), which importance was evidenced by PCA and MOFA (Figure 4 and 6). The importance of secondary metabolism in explaining natural variation between provenances was further supported by the presence of several secondary metabolites, including gibberellins and mostly phenylpropanoids, in the high confidence sPLS network generated using needles data (Figure 7). These metabolites showed a higher abundance in TAMR. The integration of the different omic layers with geoclimatic parameters highlighted, in addition, a strong direct correlation between these secondary metabolites and photosynthesis proteins, which considering the photoprotective role of phenylpropanoids present in the network supports the above-mentioned higher occurrence of photodamage in ORIA and TAMR and the need of extra photoprotection in these provenances.

The secondary metabolites in the network correlated also with the abundance of proteins involved in cellular respiration and amino acid metabolism, suggesting a metabolic rearrangement aimed at supporting the biosynthesis of secondary metabolites, especially phenylpropanoids. This rearrangement was more obvious in TAMR, accordingly with the higher abundance and importance of secondary metabolism proteins in this provenance (Figure 3C). Metabolic fluxes were redirected towards the biosynthesis of Phe, the starting point of the phenylpropanoids pathway (M. B. Pascual et al., 2016). The higher abundance of GDH1 would lead to increased levels of 2-OG in the mitochondria. 2-OG is exported from mitochondria and transported into chloroplasts through DIT1, an antiporter along with malate (Medeiros et al., 2021) found to be more abundant in TAMR. In the chloroplats, 2-OG serves as a precursor for Glu, used along with prephenate in the synthesis of Phe (M. B. Pascual et al., 2016). The synthesis of prephenate depends on the activity of prephenate dehydratase, which was between PCA a higher loading (Figure 3E). TAMR was the provenance showing the highest abundance for this enzyme. Phe is the starting point of several groups of secondary metabolites, including flavonoids or lignin (M. B. Pascual et al., 2016). The higher abundance of chloroplast metalloproteases in TAMR and their strong correlation with the abundance of some secondary metabolites suggests protein degradation at chloroplast level to generate amino acids required to support the previously exposed rearrangement involving Phe. Overall, this rearrangement implies a higher energy expenditure. It is therefore necessary to increase ATP production through cellular respiration, as it seems to happen in TAMR. The expenditure of cell resources in secondary metabolism have been described to oppose to plant growth (He et al., 2022). In concordance with this view, the lower average total growth of TAMR trees is congruent with the described metabolic rearrangement aimed at supporting the biosynthesis of photoprotective secondary metabolites like patuletin, methylsyringin or gravolenic acid (Singh et al., 2023). This metabolic rearrangement was not so obvious in ORIA, the other provenance adapted to harsh environmental conditions and that did not show an increase in secondary metabolism-related proteins (Figure 3C). In fact, average total growth of ORIA trees in the common garden was closer to that of CDVD trees (Supplemental Table 1).

Buds analysis, divided in apical and basal section, provided information on the molecular basis behind natural variation concerning development and architecture as well as on the molecular differences between bud sections, regardless of their provenance. *P. pinaster* architecture is based on a polycyclic growth pattern, i.e., in multiple shoot flushes per season. The number of flushes is related to branching degree due to the loss of apical dominance and it reduces tree value. Polycyclism is determined by environmental factors. In addition, the prevalence of specific geoclimatic features leads to the selection of specific genetic variants that also determine polycyclism, resulting in a natural variation regarding this factor. Therefore, the study of natural variation in polycyclism can shed light on such a vital process in tree development and the determination of tree architecture.

The number and type of metabolites present in different bud sections were reported to be clearly linked to the specific organogenic activity occurring in each of them (Meijón et al., 2016). Basal bud section contains axillary meristems, eventually bursting into brachyblasts, while the apical section encloses the SAM, and it is involved in its protection and maintenance. Proteomic analysis alone did not allow to separate bud sections, but its integration with previously generated metabolomics data and with environmental and growth and development features of each provenance provided new insight on bud sections biology and natural variation. The performed MOFA showed that variation at metabolome level is actually more relevant in explaining differences between apical and basal bud sections, as captured by LF2, classifying samples by section (Figure 6A, B and C). In particular, secondary metabolites characterized apical buds. These included dihydroxichalcone, a polyphenol belonging to the flavonoids family with antioxidant and ROS scavenging properties (Popovici et al., 2010). Flavonoids are also photoprotective compounds. Their accumulation in apical buds protects SAM cells from sunlight radiation damage. UV-B radiation is particularly damaging among sunlight spectrum. UV-B has a well-known phototoxic effect due to its high energy that can damage proteins and DNA (Jansen et al., 1998). Photoprotective compounds like flavonoids are a main barrier against UV-B damaging effects. UV-B sensing relies on the UV-B-specific receptor UVR8, essential for photomorphogenic responses and UV-B acclimation (Jesús Pascual et al., 2017). UVR8 was more abundant in apical buds and was present in buds high-confidence sPLS network (Figure 8A). It was connected in direct correlation to several metabolites, including propylmalic acid and several unidentified metabolites. This could be consistent with the role of UVR8 in the regulation of secondary metabolism (Rai et al., 2021). UVR8 perceives UV-B light and monomerizes into active monomers that translocate to the nucleus, where it activates the transcription of the transcription factor HY5, responsible for the expression of, among others, genes related to flavonoids biosynthesis (Stracke et al., 2010). The importance of secondary metabolism in apical buds had been previously described in the context of *P. pinaster* natural variation (Meijón et al., 2016). In this case, the integration with proteomics data added knowledge on the signaling pathway linking the accumulation of some metabolites with, in this case, environmental UV-B radiation levels.

Phytohormones are crucial in meristem maintenance and regulation in response to environmental cues, auxins being the most widely known and studied (Wang and Jiao, 2018).

Stresses decrease the number and/or size of lateral organs produced by plants, developing from axillary buds. Integration of stress signals occurs via changes in the levels and ratios between phytohormones like auxins, strigolactones or jasmonic acid, interacting with each other. Phytohormone crosstalk may modify SAM activity and change tree development and architecture. In our analysis, we identified changes in lipid metabolism related to oxylipins (Figure 4), a wide group of lipophilic compounds that include jasmonates and are involved in signaling and regulation in many defense and developmental pathways (Wasternack and Hause, 2013). The relative levels of these metabolites are a distinct characteristic of each plant species and determines the ability to adapt to different stimuli. The oxylipin pathway consists of two major branches, dependent on OAS and hydroperoxide lyase (HPL) activities, responsible for the production of jasmonates and aldehydes respectively (Farmer and Goossens, 2019). AOS was included in buds high-confidence sPLS network and showed a higher abundance in ORIA apical bud sections (Figure 4E, Figure 8A), pointing to section and provenance differences associated to jasmonates. Jasmonates have a role in a number of plant development events and in modulating them in response to the environment, regulating plant adaptation. Jasmonic acid treatments inhibit leaf expansion and hypocotyl growth (Huang et al., 2017). Overall, it has an inhibitory effect on plant growth, being accumulated especially under stress conditions, leading to a concentration of cell resources into stress response and defense rather than into growing (Major et al., 2017; Sohn et al., 2022; Zhou and Memelink, 2016). This could explain at least partially the lower total growth of ORIA trees in the common garden. UV radiation and drought are among the environmental factors stimulating jasmonates production, which would be in concordance with the higher abundance of AOS found in apical buds in general and the need of protecting the SAM from UV radiation. Within provenances, ORIA showed a higher abundance of AOS, while its level was in general lower in basal buds regardless of their provenance.

Regarding the differences between provenances, protein metabolism and vesicle trafficking highlighted as main differential processes with CDVD buds being the more different ones. This was not surprising considering CDVD is an Atlantic provenance, while ORIA and TAMR fall within the group of Mediterranean provenances (Meijón et al., 2016). Protein homeostasis and vesicle trafficking were identified as main differential processes between provenances (Figure 4C, D and E, Figure 8B). Protein homeostasis is crucial for the activity of meristems, including both its maintenance and cellular differentiation and development processes, which imply the controlled execution of specific gene expression programs as a result of the integration of both internal and external stimuli. Protein homeostasis depends on vesicle trafficking, targeting newly synthesized protein either for secretion or for degradation. Our analysis uncovered variation between provenances regarding vesicle trafficking and highlighted the importance of RAB GTPases, especially in CDVD. RABs play a significant role in plant development, regulating vesicle trafficking, cytokinesis, autophagy, and biotic and abiotic stress responses (Tripathy et al., 2021). RABs are regarded as the main organizers of the plant endomembrane network along with SNARE proteins (Nebenführ, 2002; Rodriguez-Furlan et al., 2023). Members of the RabA/Rab11 clade regulate trafficking of both secretory and endocytic vesicles through trans-Golgi network compartments. The presence of these proteins is an important determinant in cell and organ polarity, determining polarized secretion and directional fluxes that are crucial in plant growth and development as they specify cell identities and tissue and overall plant architecture, like auxin polar transport, which determines apical dominance (Yang et al., 2020).

In summary, the performed analysis evidenced that the results obtained from needles were more informative as they provided a more rich and meaningful knowledge on the biological differences between the three analyzed populations. Moreover, the integration of different layers of information, including proteomics, metabolomics, geoclimatic data and tree growing and development data, allowed a deeper and synergistic characterization of the species natural variation, providing comprehensive knowledge on the association between molecular and phenotypic information, and the influence the environment has over it.

## ACKNOWLEDGEMENTS

This paper was an output from projects financed by INIA (RTA 2010-00120-C32-01), FEDER (RTA 2013-00048-C03-02) and the Spanish Ministry of Science, Innovation and Universities (MCI-21-PID2020-113896GB-I00). JP was supported by the Juan de la Cierva Incorporación programme (IJC-2019-040330-I).

## SUPPLEMENTAL TABLE LEGENDS

**Supplemental Table 1.** Growth parameters of the three provenances used in the manuscript grown in a common garden and geoclimatic parameters at their original locations. DNI, Direct Normal Irradiance; Med, Mediterranean; Atl, Atlantic; Ap, apical; Bs, basal; Cdvd, Cadavedo; Tamr, Tamrabta.

**Supplemental Table 2.** Peaks obtained after UPLC-MS analysis of needle polar metabolites. Peaks were aligned with mzMine 2.10 avoiding redundancies between positive (P) and negative (N) modes. This table shows peak ID, detected adducts, m/z (neutral), retention time (RT) and normalized peak areas for each analyzed sample. (a) Peaks unequivocally identified (their identification was defined after comparison to our compund library or by comparison of the MS/MS to online databases). (b) Tentatively assigned peaks after comparing their accurate mass to reference compound databases. (c) Unidentified peaks. Delta ppm and compound exact mass are provided. Annotation source, molecular formula, and accessions of KEGG, FooDB and other databases are provided for all identified/assigned compounds. Assignations/identifications colors indicate their confidence (green: high, yellow: medium, red: low).

**Supplemental Table 3.** Peaks obtained after UPLC-MS analysis of bud sections polar metabolites. Peaks were aligned with mzMine 2.10 avoiding redundancies between positive (P) and negative (N) modes. This table shows peak ID, detected adducts, m/z (neutral), retention time (RT) and normalized peak areas for each analyzed sample. (a) Peaks unequivocally identified (their identification was defined after comparison to our compund library or by comparison of the MS/MS to online databases). (b) Tentatively assigned peaks after comparing their accurate mass to reference compound databases. (c) Unidentified peaks. Delta ppm and compound exact mass are provided. Annotation source, molecular formula, and accessions of KEGG, FooDB and other databases are provided for all identified/assigned compounds. Assignations/identifications colors indicate their confidence (green: high, yellow: medium, red: low).

**Supplemental Table 4.** Proteins identified in each type of sample according to SEQUEST protein search (Exp. q-value, Sum PEP Score, Coverage, number of Peptides, PSMs, Unique Peptides, Razor Peptides, number of amino acids and Protein Groups, molecular weight, isoelectric point and SEQUEST Score). Quantitative values are expressed as mean ± SD of three biological replicates. Univariate analysis results (ANOVA p-value and q-value, and TukeyHSD p-values for all paired tissue comparisons) are shown. Annotation according to Sma3s (Protein Symbol and Protein Description) and MapMan functional categories (Bnm and Bin Name) are included for each protein. Ap, apical; Bs, basal; Nd, needle; Cdvd, Cadavedo; Tamr, Tamrabta.

**Supplemental Table 5.** Results of Principal Component Analysis performed with needle and bud samples. (a) Sample Scores, (b) Protein Loadings and (c) Explained Variance. Ap, apical; Bs, basal; Nd, needle; Cdvd, Cadavedo; Tamr, Tamrabta.

**Supplemental Table 6.** Proteins identified in needles from each analysed provenance according to SEQUEST protein search (Exp. q-value, Sum PEP Score, Coverage, number of Peptides, PSMs, Unique Peptides, Razor Peptides, number of amino acids and Protein Groups, molecular weight, isoelectric point and SEQUEST Score). Quantitative values are expressed as mean ± SD of three biological replicates. Univariate analysis results (ANOVA p-value and q-value, and TukeyHSD p-values for all paired comparisons) are shown. Annotation according to Sma3s (Protein Symbol and Protein Description) and MapMan functional categories (Bin and Bin Name) are included for each protein. Nd, needle; Cdvd, Cadavedo; Tamr, Tamrabta.

**Supplemental Table 7.** Results of Principal Component Analysis performed with needle samples. (a) Sample Scores, (b) Protein Loadings and (c) Explained Variance. Nd, needle; Cdvd, Cadavedo; Tamr, Tamrabta.

**Supplemental Table 8.** Proteins identified in bud samples from each analyzed provenance according to SEQUEST protein search (Exp. q-value, Sum PEP Score, Coverage, number of Peptides, PSMs, Unique Peptides, Razor Peptides, number of amino acids and Protein Groups, molecular weight, isoelectric point and SEQUEST Score). Quantitative values are expressed as mean ± SD of three biological replicates. Univariate analysis results (ANOVA p-value and q-value, and TukeyHSD p-values for all paired tissue comparisons) are shown. Annotation according to Sma3s (Protein Symbol and Protein Description) and MapMan functional categories (Bnm and Bin Name) are included for each protein. Ap, apical; Bs, basal; Nd, needle; Cdvd, Cadavedo; Tamr, Tamrabta.

**Supplemental Table 9.** Results of Principal Component Analysis performed with bud samples. (a) Sample Scores, (b) Protein Loadings and (c) Explained Variance. Nd, needle; Cdvd, Cadavedo; Tamr, Tamrabta.

**Supplemental Table 10.** Multi-Omic Factor Analysis (MOFA) results. (a) % of variance explained at each omic level in needles MOFA and (b) divided by latent factor (LF). (c) % of variance explained at each omic level in buds MOFA and (b) divided by LF.

**Supplemental Table 11.** Gene Set Enrichment Analysis of MOFA features in LF1 and 2 in (a) needles and (b) buds according to MapMan functional classification. pval.adj, FDR.adjusted p-value; nfeatures, number of features in each category; sign, correlation value sign.

## Footnotes

1 https://github.com/Valledor/pRocessomics

